# Molecular mechanics of smooth muscle contraction and relaxation modulated by caldesmon

**DOI:** 10.64898/2026.03.23.713758

**Authors:** Matheus L. C. Schultz, Linda Kachmar, Christie Liu, Andy Bai, Sean Fletcher, Anne-Marie Lauzon

## Abstract

Smooth muscle (SM) contraction is well known to be regulated by the reversible phosphorylation of the myosin regulatory light chain. However, SM force generation and relaxation are often uncoupled from myosin phosphorylation levels (e.g. the latch-state), indicating that additional regulatory mechanisms must be at play. The precise effects of the actin binding protein caldesmon (CaD) on SM force production and relaxation remain ambiguous, largely due to contradictory findings in experiments performed at the tissue level. To date, there are no studies that have measured the effects of CaD on force and relaxation at the molecular level. Here, we use a laser-trap assay to measure the force produced by SM myosin molecules in the presence and absence of CaD. Measurements were performed before and during myosin dephosphorylation, thus simulating SM contraction and relaxation in-vitro. We demonstrate that CaD inhibits force generation, most likely through competitive inhibition of actomyosin binding while simultaneously introducing a resistive load via tethering of actin and myosin. We also establish CaD as a potentiator of relaxation, increasing force decay rate during myosin dephosphorylation. Finally, we show that CaD directly modulates the dependence of myosin-actin mechanics on myosin phosphorylation levels. These findings refine our understanding of SM regulation, highlighting CaD not merely as a passive structural stabilizer, but as a critical regulatory component of force development and relaxation. Ultimately, understanding these mechanical functions offers new perspectives on pathophysiologies involving SM, such as asthma, hypertension, and gastrointestinal disorders, potentially guiding targeted therapeutic strategies.

**SIGNIFICANCE STATEMENT:** Smooth muscle (SM) is responsible for controlling the internal diameter of blood vessels and viscera. Understanding the precise regulation of SM relaxation by actin-binding proteins remains a fundamental lacuna in physiology. Using a molecular mechanics chamber to manipulate the biochemical milieu during active measurements, we demonstrate, for the first time at the molecular level, that caldesmon (CaD) acts as a mechanical modulator that inhibits force generation and accelerates relaxation of SM myosin ensembles. Our results provide a molecular basis for resolving previous contradictory findings reported in tissue-level experiments. Ultimately, understanding the role of contractile and regulatory proteins of SM will provide the basis for understanding SM disorders, such as hypertension and asthma, and guide the development of targeted therapeutic strategies.

## INTRODUCTION

Smooth muscle (SM) is responsible for the contraction of all hollow organs in the body, except for the heart, and its dysregulation can contribute to conditions such as asthma, hypertension, and gastrointestinal pathologies. Contraction is initiated by an increase in intracellular Ca^2+^, which binds to calmodulin and together activate myosin light chain kinase (MLCK). MLCK phosphorylates the 20 kDa regulatory light chain (LC_20_) of SM myosin, enabling myosin-actin interactions that generate force through ATP hydrolysis. Relaxation occurs via LC_20_ dephosphorylation by myosin light chain phosphatase (MLCP). Despite being well accepted, the mechanism described above is not sufficient to fully explain SM contractile regulation, because force generation and relaxation are often uncoupled from myosin phosphorylation levels (1–5).

Caldesmon (CaD) is an actin and myosin binding protein thought to be involved in the regulation of SM contraction. Evidence for this role comes mostly from in-vitro studies showing that CaD inhibits the actomyosin ATPase activity by hindering the binding of myosin to the actin filament (6, 7). CaD also tethers actin and myosin filaments together, which is thought to help maintain the structural integrity of the contractile apparatus (8). This is supported by studies that showed that CaD enhances the binding of myosin to actin (6, 7, 9–11) and that it distributes in clusters in SM, where actin and myosin filaments intercross (12). Additionally, in non-muscle cells, CaD has been shown to ensure regular spacing of myosin heads within the cytoskeleton (13).

The effects of CaD on SM force generation and relaxation have been studied exclusively at the tissue level, where it has been reported that CaD increases the rate of SM relaxation (14–17). However, the effects of CaD on force development remain unclear, since different methodologies have yielded conflicting results. For instance, while exogenous addition of CaD suppressed force (18, 19), partial genetic knockout (KO) caused force to increase (17). In contrast, full KO or knockdown often paradoxically caused force reduction (15–17, 20). Tissue models face significant limitations as the complete genetic loss of CaD is perinatally lethal (16), and partial depletion often triggers compensatory changes in protein expression and post-translational modifications that can lead to misinterpretation of the contractile outcome (14, 15). In terms of its effects on actomyosin molecular mechanics, CaD is known to reduce actin filament propulsion velocity (*v*_*avg*_) in the in-vitro motility assay (IVMA) (10, 11, 21, 22) likely by crosslinking the actin filaments to the myosin molecules (23, 24). However, no molecular mechanics studies have explored the effects of CaD on force generation and relaxation of SM myosin molecules.

We recently developed assay chambers that enable alterations of experimental conditions during molecular mechanics measurements (25), allowing the simulation of SM relaxation in-vitro. In this study, we used this device to measure the force produced by SM myosin molecules in the presence and absence of CaD, before and during myosin dephosphorylation, thus simulating SM contraction and relaxation in-vitro. We showed, for the first time at the molecular level, that CaD inhibits force production, and accelerates relaxation during myosin deactivation.

## RESULTS

To assess how CaD modulates force-generation, force maintenance and relaxation during myosin dephosphorylation, we allowed a group of myosin molecules to pull on an actin filament attached to a laser-trapped microsphere until force reached a plateau. At this steady state, MLCP was injected into the system to initiate myosin deactivation, resulting in force decay (Fig. 1).

**Figure 1:**
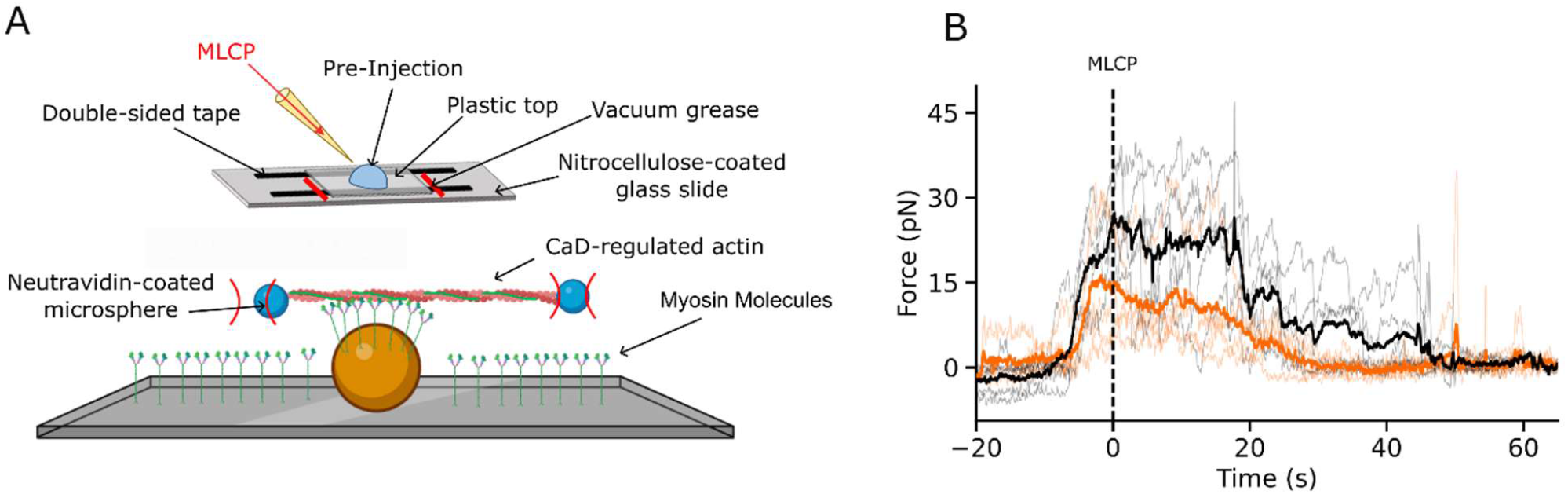
Experimental setup. (A) Schematic representation of the laser trap assay and the flow-through chamber used for MLCP injections. (B) Individual force traces (faint lines) and their corresponding averages (bold lines) recorded in the absence (black) or presence (orange) of 250nM CaD. Traces were aligned to the MLCP injection time (t = 0 s, dashed vertical line).

### CaD reduces maximum force

To characterize how CaD alters force development kinetics before MLCP injection, the rising portion of force traces were ensemble-averaged and fitted with a sigmoidal function yielding values for the force-rise rate (*k*_*rise*_) and force plateau (*F*_max_) (Fig. 2A). To assess the distribution of these parameters, we performed an exact bootstrap analysis by generating all unique combinations with replacement of the force events. Each combination was ensemble-averaged and fitted to the sigmoidal model, yielding a complete distribution of kinetic parameters (Figs. 2B and C; Table 1).

**Figure 2:**
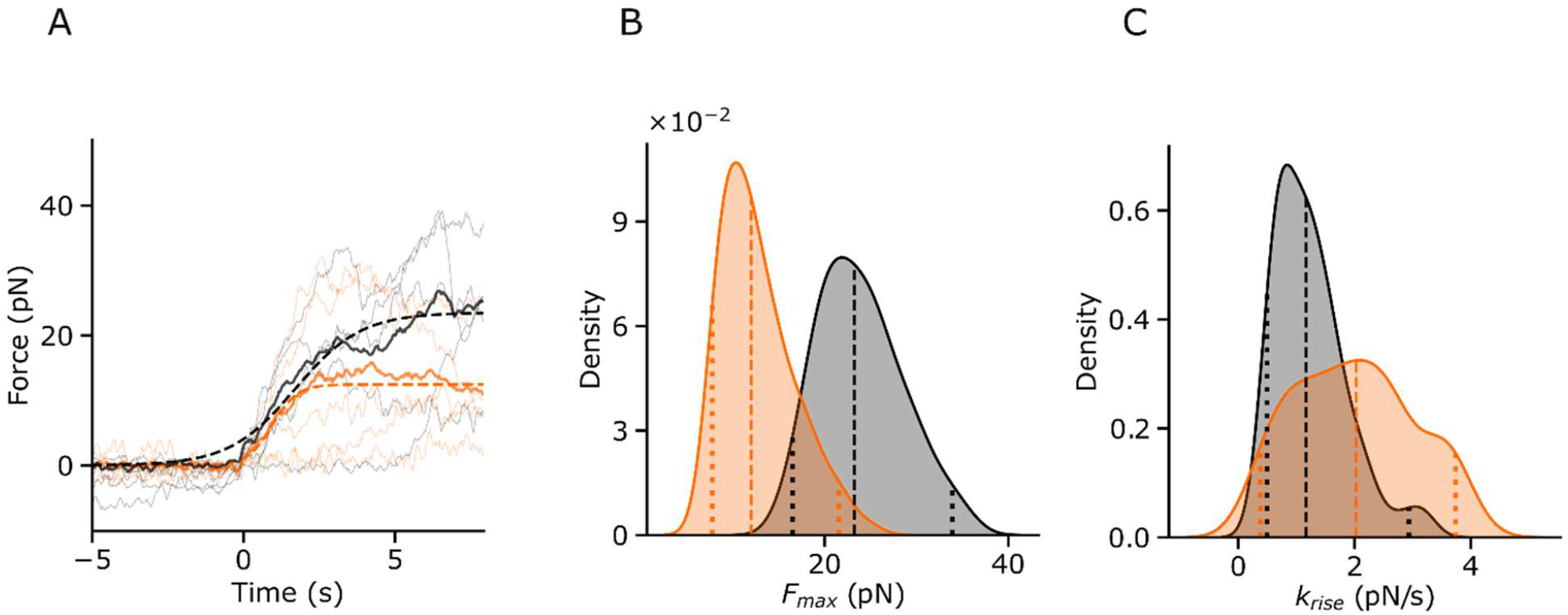
Analysis of force increase kinetics. Data in the presence of 250nM CaD is shown in orange; data in its absence is shown in grey. (A) Sigmoidal functions (dashed lines) fitted to the ensemble-averaged force traces (bold solid lines). Individual traces (n = 5 per condition) are shown as faint lines. (B and C) Kernel density estimation of *F*_*max*_ and *k*_*rise*_ derived from sigmoidal fits to all unique combinations with replacement of the collected force events (n = 126 combinations). Vertical dashed lines indicate the median. Vertical dotted lines indicate the 95% CI.

**Table 1:**
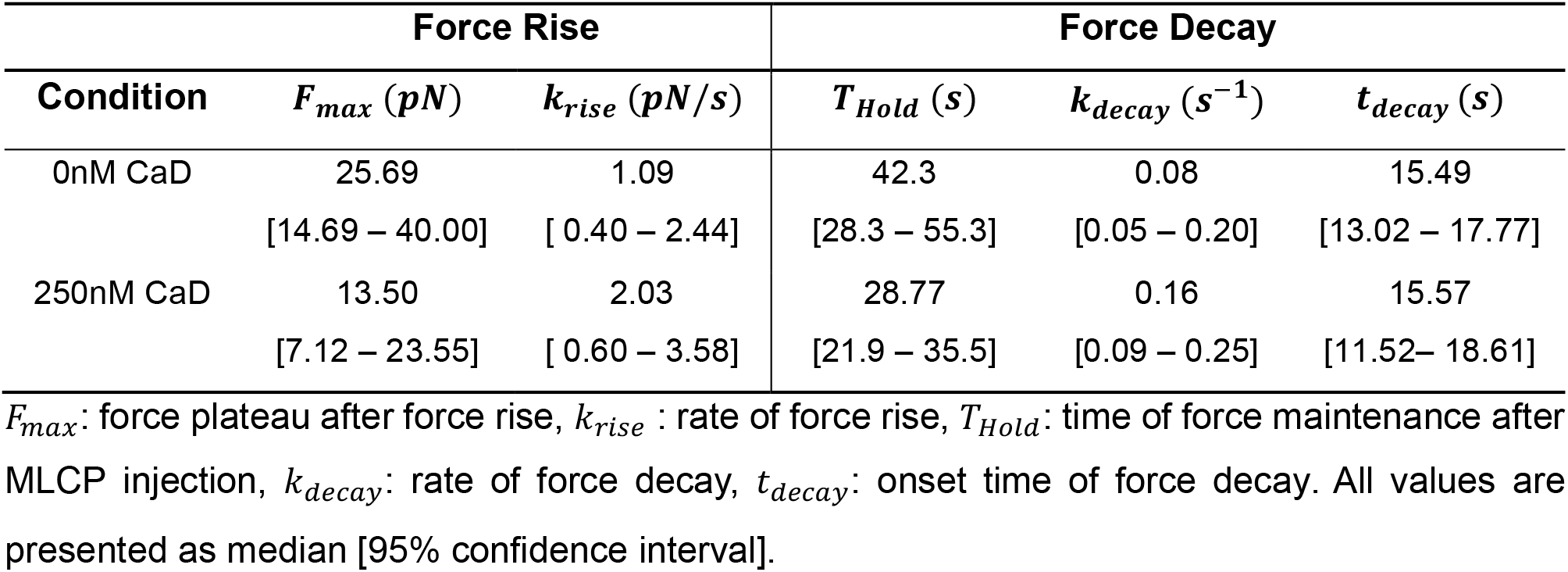
Force rise and decay kinetics parameters.

The resulting parameter distributions revealed that the median *F*_*max*_ was substantially suppressed by CaD from a control value of 23.25 pN to 12.02 pN, with ∼97% of the combinations exhibiting a lower *F*_*max*_ with CaD (Cliff’s *δ* = -0.93). In contrast, although CaD slightly increased the median *k*_*rise*_ from a control value of 1.17pN/s to 2.03 pN/s, the distributions exhibited substantial overlap, with ∼72% of combinations yielding a higher *k*_*rise*_ with CaD (Cliff’s *δ* = 0.56). Together, these results demonstrate that CaD limits force generation capacity without robust alterations in the rate of force development.

### CaD decreases the time of force maintenance by accelerating force decay

To assess how CaD affects force maintenance during myosin dephosphorylation, we quantified the duration of force events following MLCP injection (*T*_*Hold*_) (Fig. 3A & see methods). Again, we performed an exact bootstrap analysis by generating all unique combinations of the collected events with replacement and calculated the mean *T*_*Hold*_ for each combination, yielding a complete *T*_*Hold*_ distribution with and without CaD (Fig. 3B). CaD substantially reduced the median *T*_*Hold*_ from a control value of 42.87 s to 27.80 s, with 88% of combinations exhibiting a shorter *T*_*Hold*_ with CaD (Cliff’s *δ* = 0.76). Importantly, these differences were not caused by differences in the timing of injection between conditions (see supplementary material).

**Figure 3:**
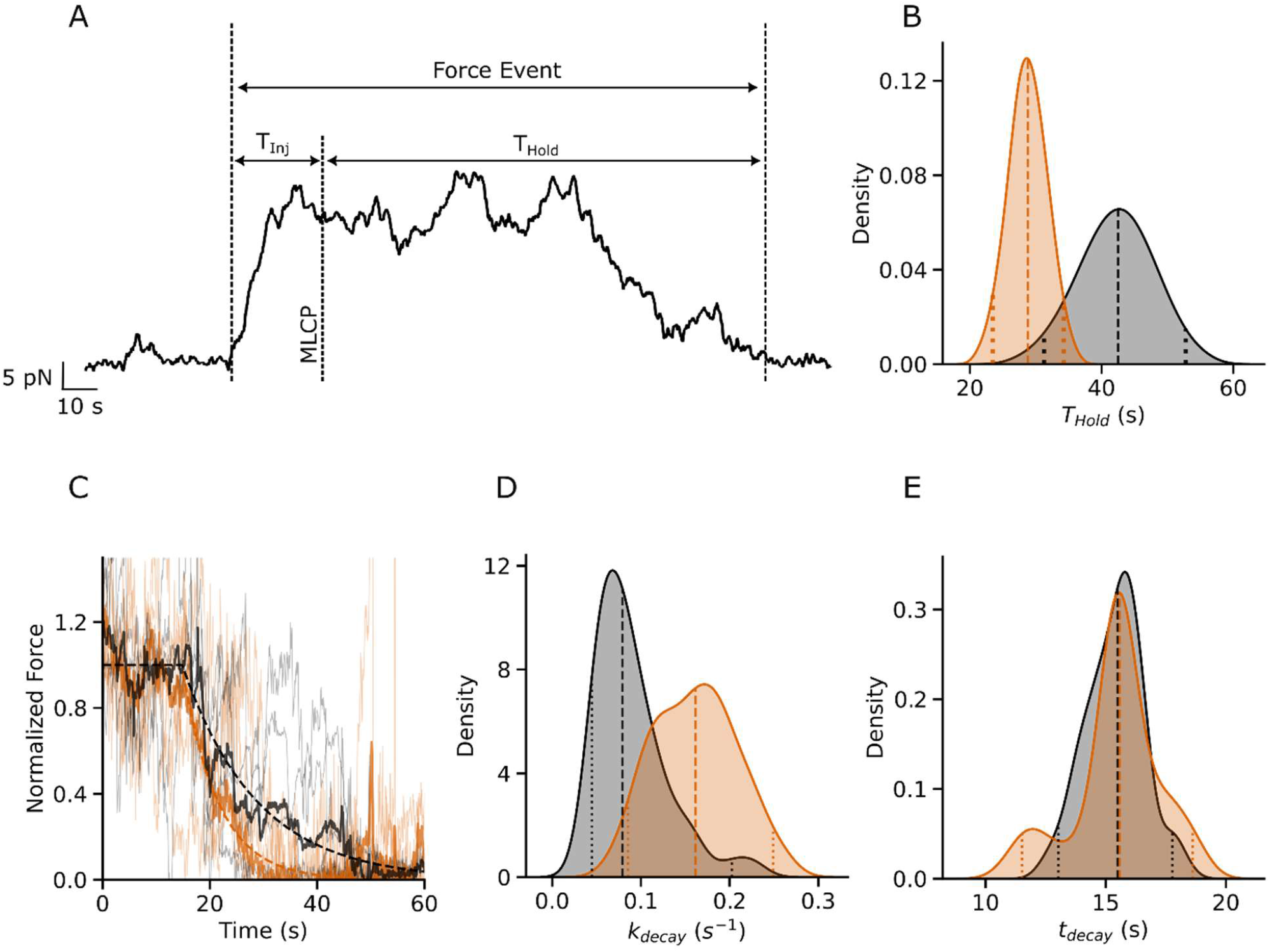
Analysis of force maintenance and force decay after MLCP injection. Data in the presence of 250nM CaD is shown in orange; data in its absence is shown in grey. (A) Representative force trace illustrating the definition of *T*_*Hold*_ and *T*_*/nj*_. (B) Kernel density estimation of average *T*_*Hold*_ calculated as the average *T*_*Hold*_ from all unique combinations with replacement of the collected force events (n = 126 combinations). (D) Shifted exponential decay functions (dashed lines) fitted to the ensemble-averaged force traces (bold solid lines). Individual traces are shown as faint lines. (E and F) Kernel density estimation of *k*_*decay*_ and *t*_*decay*_ derived from sigmoidal fits to all unique combinations with replacement of the collected force events. Vertical dashed lines indicate the median. Vertical dotted lines indicate the 95% CI.

To determine how CaD affects force decay kinetics, the decaying portion of force-events were ensemble-averaged and normalized to the steady-state plateau, defined as the mean force during the first 10 s following MLCP injection. These normalized ensemble-traces were fitted with a shifted exponential decay function, yielding parameters for the rate (*k*_*decay*_) and the onset (*t*_*decay*_) of force decay (Fig. 3C). The distribution of *k*_*decay*_ and *t*_*decay*_ were determined with the same approach described above for the force-rise analysis, yielding a distribution of decay parameters (Figs. 3D and E; Table 1). These distributions revealed a greater *k*_*decay*_ in the presence of CaD, from 0.08 s^-1^ for control to 0.16 s^-1^ with CaD, with ∼88% of combinations exhibiting a higher *k*_*decay*_ with CaD (Cliff’s *δ* = 0.77). By contrast, there was no difference in the median *t*_*decay*_ between conditions (Control: 15.5 s; CaD: 15.6 s; Cliff’s *δ* = 0.06). Together these results demonstrate that CaD accelerates relaxation rates after myosin dephosphorylation.

### CaD enhances the deceleration of actin during myosin dephosphorylation in the IVMA

To gain further insights into the effects of CaD on actomyosin molecular mechanics during myosin dephosphorylation, we measured *v*_*avg*_ and the fraction of moving filaments (*f*_*mot*_) following MLCP injection in the IVMA. As expected, both *v*_*avg*_ and *f*_*mot*_ decreased after MLCP injection both in the presence and absence of CaD (Figs. 4A and D). To characterize the kinetics of this motility decay, we performed a stochastic bootstrap analysis (2,000 iterations) of the normalized *v*_*avg*_ and *f*_*mot*_ traces to generate the parameter distributions.

**Figure 4:**
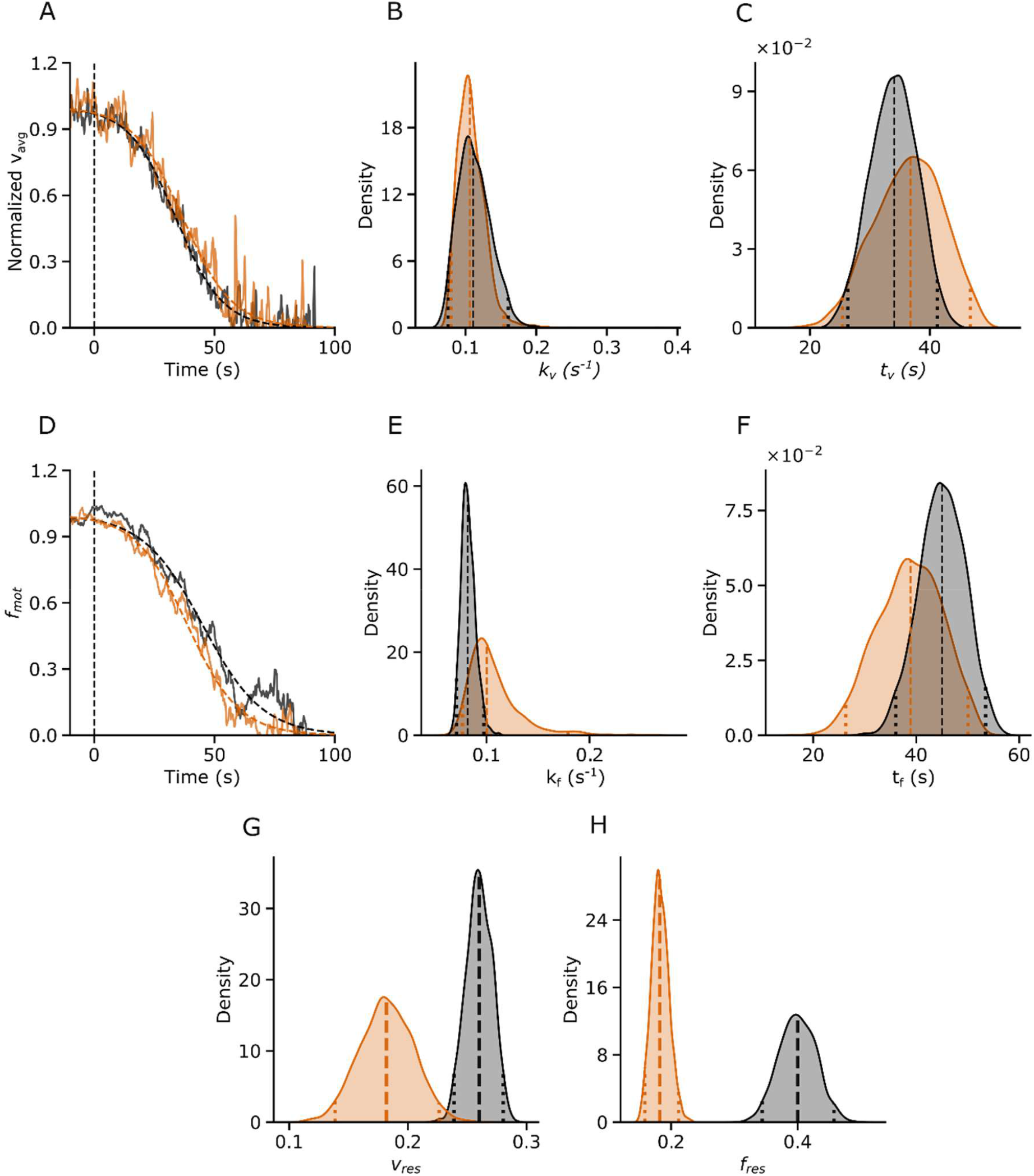
Analysis of *v*_*avg*_ and *f*_*mot*_ decay after MLCP injection in the IVMA. Data in the presence of 250nM CaD is shown in orange; data in its absence is shown in grey. (A, E) Sigmoidal functions (dashed lines) fitted to the ensemble-averaged *v*_*avg*_ (A) and *f*_*mot*_ (E) traces (bold solid lines). Individual traces (n =10) are omitted for clarity. Vertical dashed lines show the time of MLCP injection. (B, C, F, G) Kernel density estimation of the sigmoidal rate and midpoint derived from 2000 bootstrap iterations of *v*_*avg*_ (B and C) and *f*_*mot*_ (F and G) traces. (D and H) Kernel density estimation of the averages of *v*_*res*_ and *f*_*res*_ from 2000 bootstrap samples of the *v*_*avg*_ and *f*_*mot*_ traces. Vertical dashed lines indicate the median. Vertical dotted lines indicate the 95% CI.

Ensemble-averaged samples were fitted with a sigmoidal function yielding the decay rate (*k*_*v*_ for *v*_*avg*_ or *k*_*f*_ for *f*_*mot*_) and the sigmoidal midpoint (*t*_*v*_ for *v*_*avg*_ or *t*_*f*_ for *f*_*mot*_). For *v*_*avg*_, the presence of CaD had marginal to no effects on the sigmoidal parameters. The median *k*_*v*_ was similar between conditions (*∼ 0*.*11* s^-1^), with only ∼55% of bootstrap samples having a slower *k*_*v*_ with CaD (Cliff’s *δ* = -0.11). Similarly, CaD shifted the median *t*_*v*_ from 34.05 s for control to 36.77 s, yielding slightly higher values in just ∼64% of the CaD samples (Cliff’s *δ* = 0.29) (Figs. 4B and C; Table 2).

**Table 2:** *v*_*avg*_ and *f*_*mot*_ decay parameters in the IVMA.

| Condition | $v_{avg}$ | | | $f_{mot}$ | | |
| --- | --- | --- | --- | --- | --- | --- |
| | $k_v (s^{-1})$ | $t_v (s)$ | $v_{res}$ | $k_f (s^{-1})$ | $t_f (s)$ | $f_{res}$ |
| 0nM CaD | 0.11 | 34.05 | 0.26 | 0.08 | 45.02 | 0.40 |
|  | [0.08 – 0.15] | [26.30 – 41.24] | [0.24 – 0.28] | [0.07 – 0.10] | [36.02 – 53.47] | [0.34 – 0.46] |
| 250nM CaD | 0.11 | 36.77 | 0.18 | 0.10 | 38.90 | 0.18 |
|  | [0.08 – 0.16] | [25.41 – 46.77] | [0.14 – 0.23] | [0.08 – 0.18] | [26.34 – 50.08] | [0.16 – 0.21] |

$k_v$ and $k_f$ : sigmoidal rates, $t_v$ and $t_f$ sigmoidal midpoints, $v_{res}$ and $f_{res}$ : normalized residual values for $v_{avg}$ and $f_{mot}$ , respectively. All values are presented as median [95% confidence interval].

In contrast, CaD exerted a more pronounced effect on the *f*_*mot*_ decay kinetics. The median *k*_*f*_ increased from 0.082 s^-1^ for control to 0.100 s^-1^ with CaD, with ∼88.4% of bootstrap samples having a higher *k*_*f*_ with CaD (Cliff’s *δ* = 0.77). Furthermore, *t*_*f*_ decreased from 45.02 s for control to 38.90 s with CaD, with ∼78% of bootstrap samples having a lower *t*_*f*_ with CaD (Cliff’s *δ* = -0.57) (Figs. 4E and F; Table 2).

Next, we quantified the average normalized residual velocity (*v*_*res*_) and motile fraction (*f*_*res*_) after myosin dephosphorylation using the same stochastic bootstrapping approach. The addition of CaD caused a substantial decrease in *v*_*res*_ from a median of 0.26 for control to 0.18 with CaD (Fig. 4G) and in *f*_*res*_ from a median of 0.40 for control to 0.18 with CaD (Fig 4H), yielding a maximum effect size of Cliff’s *δ* = 1 for both parameters.

### CaD changes the relationship between myosin phosphorylation levels and *v*_*avg*_ or *f*_*mot*_

We measured *v*_*avg*_ and *f*_*mot*_ at varying myosin phosphorylation levels to determine if CaD alters the mechanical sensitivity to myosin phosphorylation (Fig. 5). The control relationship (without CaD) exhibited a saturating dependence on phosphorylation, but the presence of CaD qualitatively altered this relationship, resulting in a quasilinear relationship for *v*_*avg*_ with no observable plateau. Statistical model selection favored a linear fit over a rectangular hyperbola for the CaD group (*Δ*AIC = 0.25; Supplemental Table 1). The specificity of the effect of CaD was confirmed using bovine serum albumin (BSA) as a negative control. Unlike CaD, BSA did not alter the phosphorylation-dependence curves, which remained statistically indistinguishable from the control fits (F-test; p = 0.95 for *v*_*avg*_, p = 0.49 for *f*_*mot*_). Best-fit parameters of the fitted models are shown in Table 3. Together, these results show that CaD reduces the sensitivity of *v*_*avg*_ and *f*_*mot*_ to myosin phosphorylation.

**Figure 5:**
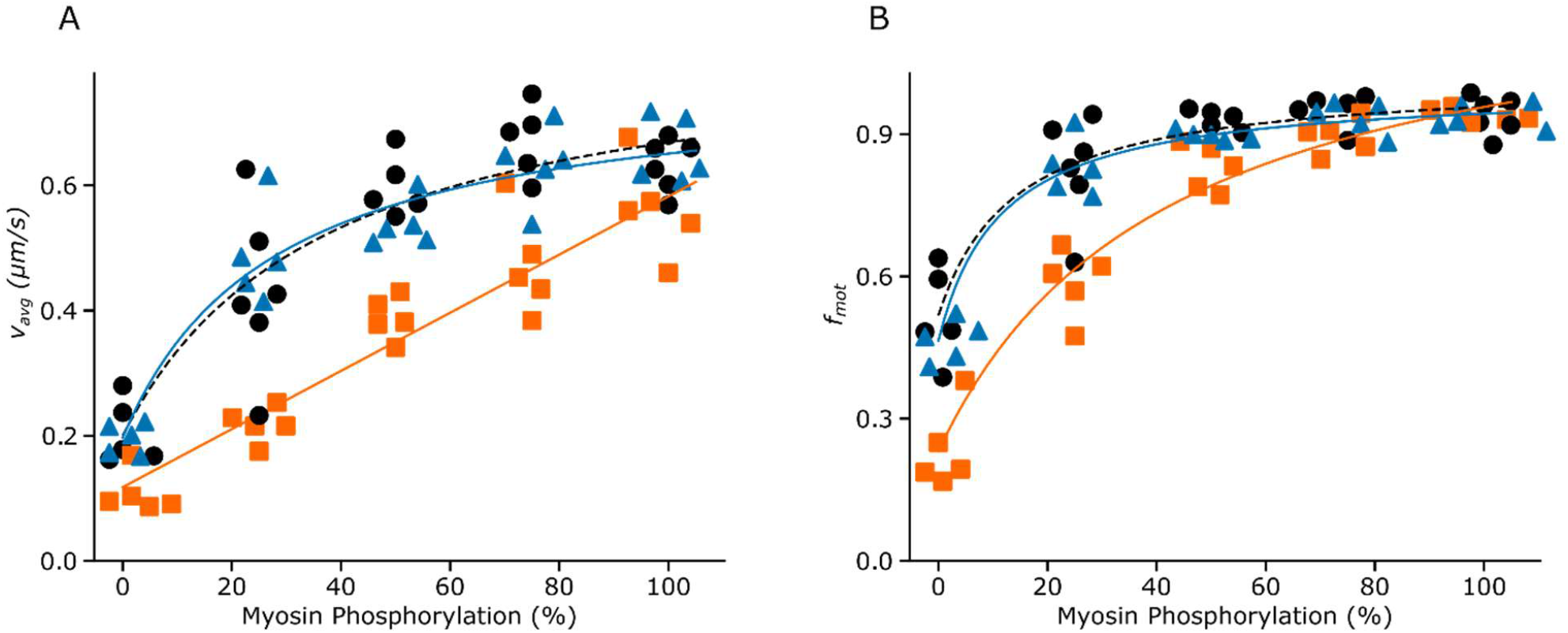
*v*_*avg*_ and *f*_*mot*_ dependence on myosin phosphorylation levels. (A) *v*_*avg*_ vs myosin phosphorylation levels. (B) *f*_*mot*_ vs myosin phosphorylation levels. Experiments were done in the absence (black) or the presence of 250nM CaD (orange) or BSA (blue). Overlayed lines show rectangular hyperbola fits except for the dataset of *v*_*avg*_ in the presence of CaD, which was fitted with a linear model. Data points are horizontally jittered to prevent overlap; experiments were performed at discrete phosphorylation levels of 0, 25, 50, 75, and 100% phosphorylation.

**Table 3:** Parameters obtained from the mixture motility assay.

| Metric | Condition | Model | $y_{max}$ or $s$ | $y_0$ | $k_m$ | $R^2$ |
| --- | --- | --- | --- | --- | --- | --- |
| $v_{avg}$<br>$\mu m/s$ | Control | Hyperbola | $0.84 \pm 0.1$ | $0.20 \pm 0.04$ | $37 \pm 17$ | 0.83 |
| | 250nM CaD | Linear | $4.6 \pm 0.3 \times 10^{-3}$ | $0.12 \pm 0.02$ | --- | 0.90 |
| | 250nM BSA | Hyperbola | $0.78 \pm 0.06$ | $0.20 \pm 0.02$ | $28 \pm 9$ | 0.91 |
| $f_{mot}$ | Control | Hyperbola | $1.00 \pm 0.05$ | $0.52 \pm 0.03$ | $12 \pm 6$ | 0.85 |
| | 250nM CaD | Hyperbola | $1.00 \pm 0.07$ | $0.23 \pm 0.04$ | $18 \pm 6$ | 0.92 |
| | 250nM BSA | Hyperbola | $1.00 \pm 0.03$ | $0.46 \pm 0.02$ | $12 \pm 3$ | 0.96 |

A rectangular hyperbola or a linear model was fitted to the data of both $v_{avg}$ and $f_{mot}$ vs phosphorylation levels obtained from mixture motility assays. The rectangular hyperbola fits yielded values for an upper asymptote ( $y_{max}$ ), a y-axis intercept ( $y_0$ ), and the phosphorylation level at 50% maximum $v_{avg}$ or $f_{mot}$ ( $k_m$ ). In addition to the y-axis intercept, the linear model also yielded a slope value ( $s$ ). The goodness of fit is presented by the $R^2$ values. All parameters are presented as mean ± sem.

## DISCUSSION

This study provides new insight into the role of CaD as a mechanical modulator of SM contraction. Although CaD is well-characterized as an inhibitor of actomyosin ATPase (6, 7, 26), its specific effects on force generation and relaxation remain unclear. We showed that CaD functions not only as a molecular break on force development, but it also accelerates relaxation during myosin dephosphorylation. Specifically, the main findings of this study, performed in vitro with an ensemble of SM myosin molecules, are: 1) CaD reduces force production, 2) CaD reduces the time of force maintenance during dephosphorylation, 3) CaD accelerates the rate of relaxation, and 4) CaD alters the dependence of myosin-actin mechanics on myosin activation levels.

### Effects of CaD on Force Production

Integrating CaD in the laser-trap assay substantially reduced *F*_*max*_. This result is in agreement with tissue studies reporting a decrease in SM force generation upon addition of exogenous CaD (18, 19). One possible explanation for this CaD-induced force reduction is its ability to physically block myosin from binding to actin filaments. Several studies have shown that the actin-binding domain of CaD competes with myosin for actin attachment (6, 7, 26–28). This competitive inhibition would result in fewer myosin heads generating force.

Alternatively, CaD could tether myosin to actin, creating a passive load that opposes the force generated by cycling cross-bridges. However, such a load appears insufficient to fully account for our observed drop in *F*_*max*_. Roman and coworkers reported that CaD increases the force required to detach a group of unphosphorylated SM myosin molecules from an actin filament by ∼0.06 pN/myosin, a value they derived by normalizing their total measured force by the number of myosin molecules in proximity to the filament (11).

Because the binding affinity of CaD to myosin is independent of the myosin phosphorylation state (29), a drag-force of similar amplitude would be expected in our assay. In contrast, when applying the same normalization method to our *F*_*max*_ measurements (see Supplementary Material) we observed a force reduction of ∼0.48 pN/myosin in the presence of CaD, which represents a load ∼8-fold greater than that reported by Roman and colleagues (11). Therefore, although the observed force reduction may be a combination of both the passive drag and the hindrance mechanism of CaD, the latter appears to be the dominant factor due to the small amplitude of the previously reported load (11). It remains unclear whether CaD alters the mechanics of individual myosin molecules or only affects them in groups. Resolving this question will require single-molecule measurements.

Interestingly most tissue-based studies report a decrease in force upon removal of CaD, which contrasts with our findings (15, 16, 20). However, Pütz et al. demonstrated that this decrease stems primarily form CaD’s structural role in maintaining the integrity of the contractile apparatus (16). They observed that force reduction in CaD KO samples is more pronounced in permeabilized tissues, where the loss of CaD leads to myosin depletion. While a force decrease also occurs in non-permeabilized KO tissue, the effect only becomes prominent under high stretch, again disturbing the muscle integrity (16). To our knowledge, our study is the first to address the effect of CaD on force and relaxation at the molecular level. This approach allowed us to decouple CaD’s structural role from direct regulation of myosin-actin mechanics, as the actomyosin structural integrity of our in-vitro assay was independent of CaD.

The mild increase in *k*_*rise*_ in the presence of CaD was unexpected, as it contrasts with the well-documented ability of CaD to decrease *v*_*avg*_ in the IVMA (Fig. S2) (10, 11, 21, 22). This discrepancy suggests that CaD may operate differently under loaded conditions. This is consistent with Smolock and colleagues, who observed a slight decrease in isotonic shortening velocity in CaD-knockdown tissue (20). However, future experiments with controlled loads are necessary to confirm this load-dependent mechanism.

### Effects of CaD on Force Decay

In addition to lowering *F*_*max*_, we found that CaD shortens *T*_*Hold*_. Shifted-exponential fits of the normalized force relaxation traces revealed that this effect stems from a higher rate of force decay following MLCP injection. This observation aligns with several tissue-level studies reporting that CaD accelerates SM relaxation (14, 16, 17). Albrecht et al. previously proposed that CaD increases relaxation rates by inhibiting cooperative binding of dephosphorylated myosin (14), a hypothesis supported by findings that CaD inhibits the cooperative activation of thin filaments by rigor myosin heads (6). However, the cooperative recruitment of dephosphorylated myosin requires tropomyosin (30), which was absent in our experiments. Consequently, the accelerated relaxation we observed cannot be attributed to thin-filament-induced cooperativity. Instead, these results suggest that CaD exerts a direct effect on the actomyosin interaction itself.

We propose that this accelerated force decay is driven by a CaD-induced desensitization of myosin-actin mechanics to myosin phosphorylation levels. Previous studies in skinned tissue have demonstrated that CaD alters the force-phosphorylation relationship, requiring higher levels of myosin phosphorylation to maintain the same amount of force (18, 19, 31). Consistent with this, our IVMA results showed a similar uncoupling of *v*_*avg*_ and *f*_*mot*_ to myosin phosphorylation. This desensitization provides a direct explanation for the faster relaxation: since CaD reduces the mechanical performance at any given phosphorylation levels, the continuous dephosphorylation by MLCP leads to a more rapid collapse in the total mechanical output of the system.

### Effects of CaD on actin filament deceleration during MLCP Injection in the IVMA

In the IVMA, the addition of CaD accelerates the decay of *f*_*mot*_ following MLCP injection. This aligns with the faster relaxation observed in the laser-trap assay and likely stems from the same CaD-induced desensitization of contractile mechanics to myosin phosphorylation levels. Surprisingly, this did not translate into a faster decay of *v*_*avg*_. We suspect this discrepancy is an artifact of the *v*_*avg*_ calculation, which excludes arrested filaments. The combination of increased *k*_*f*_ and unchanged *k*_*v*_ suggests that CaD triggers abrupt halting rather than a gradual, collective deceleration during myosin dephosphorylation. Consequently, in the presence of CaD, the *v*_*avg*_ traces count fewer filaments over time. This introduces more noise into the signal, leading to an unreliable *v*_*avg*_ decay estimation.

Alternatively, filaments with an inherently faster *v*_*avg*_ may be less likely to halt abruptly. As myosin becomes dephosphorylated and slower filaments stop (and are thus excluded from the calculation), the measured *v*_*avg*_ becomes artificially inflated. This ‘survivor bias’ would result in an apparently slower *k*_*v*_ and a shift in the sigmoidal midpoint to later times, consistent with our modest observed *t*_*v*_ shift. This raises the question of whether the subset of filaments that continues to move is simply “unregulated” (i.e., lacking CaD). However, the lower *v*_*res*_ in the presence of CaD compared to control indicates that the filaments that continue moving are indeed suppressed by CaD.

Finally, it’s possible that CaD alters the rate of myosin dephosphorylation itself. This is plausible given that CaD binds directly to myosin (32, 33), and tissue-level studies have shown that the myosin-binding domain of CaD is critical for modulating SM relaxation rates (31). However, our analytical estimation of the dephosphorylation kinetics (see Supplementary Material) suggests that CaD has no such effect. Note that this estimation relies on the assumption that the mechanical output at any given phosphorylation level is independent of whether that level was reached via active dephosphorylation or controlled steady state conditions; therefore, it should be interpreted with caution. Nevertheless, it provides theoretical evidence that the observed changes in *v*_*avg*_, *f*_*mot*_, and force decay result from altered mechanics rather than shifted dephosphorylation rates.

## Conclusion

In summary, this study provides a comprehensive *in vitro* characterization of CaD as a mechanical modulator of SM contraction. For the first time at the molecular level, we demonstrated that CaD inhibits force generation, most likely through competitive inhibition of actomyosin binding while simultaneously introducing a resistive load via tethering of actin and myosin. Furthermore, our results establish CaD as a potentiator of relaxation, increasing the force decay rate during myosin dephosphorylation. These findings refine our understanding of SM regulation, highlighting CaD not merely as a passive structural stabilizer, but as an important regulatory component of force development and relaxation. Ultimately, understanding these mechanical functions of CaD may offer new perspectives on pathophysiologies involving SM, such as asthma, hypertension, and gastrointestinal disorders, potentially guiding the development of more targeted therapeutic strategies.

## METHODS

### Reagents

The following reagents were purchased from Sigma-Aldrich: ATP (A3377), BSA (A7030), CaCl_2_ (C5080), calmodulin (P2277), catalase (C40), glucose (G7528), glucose oxidase (G2133-50KU), glycerol (356352), imidazole (I202), methylcellulose (M0512), MnCl_2_ (203734), and tetramethyl rhodamine isothiocyanate (TRITC)-phalloidin (P1951).

The following reagents were purchased from Thermo Fisher Scientific: amyl acetate (A718-500), dithiothreitol (DTT; BP172), EGTA (O2783), glutaraldehyde (BP25484), KCl (P330), and MgCl_2_ (M33-500).

### Buffers

Buffer concentrations were as follows (in mM unless specified otherwise).

#### Myosin Buffer

300 KCl, 25 imidazole, 1 EGTA, 4 MgCl_2_, 15 DTT, pH 7.4.

#### Actin Buffer

25 Imidazole, 25 KCl, 4 MgCl_2_, 0.1 CaCl_2_, pH 7.4.

#### IVMA Injection Buffer

Actin buffer with 2 ATP, and 1 MnCl_2_.

#### Injection Buffer

Actin buffer with 0.2 ATP, and 1 MnCl_2_.

#### Motility Assay Buffer

Actin buffer with 0.5% methylcellulose, 2 ATP, and 1 MnCl_2_.

#### Laser Trapping Assay Buffer

Actin buffer with 0.3% methylcellulose, 0.2 ATP, 1 MnCl_2_.

### Proteins

SM myosin was purified from chicken gizzard (34). SM CaD was purified from pig stomach fundus (35). MLCK and MLCP were purified from turkey gizzard (36, 37). The MLCP preparation includes the catalytic (PP1-37kD) and target (MyPT1-67kD) subunits. CaD, MLCK, and MLCP were kindly provided by Dr. A. Sobieszek, Austrian Academy of Sciences, Vienna, Austria.

SM myosin (5 mg/ml) was phosphorylated by incubation with CaCl_2_ (6 mM), calmodulin (3 μM), MLCK (0.075 μM), MgCl_2_ (10 mM), and MgATP (5 mM) for 20 min at room temperature. The samples were then stored in 50% glycerol at −20°C and used within 72h. Immediately before the assays, phosphorylated SM myosin (500 μg/mL) was ultracentrifuged (Beckman XPN-90; 42.2 Ti rotor; 31 min; 4°C; 42,000 rpm) in myosin buffer with filamentous actin (100 μg/mL) and ATP (1mM), to remove non-functional myosin molecules.

Biotinylated-TRITC-labelled actin filaments were prepared by polymerizing actin filaments with a mixture of actin monomers where 25% of monomers were biotinylated. The polymerized actin was labelled with TRITC-phalloidin. Actin biotinylation did not affect the regulation of *v*_*avg*_ or *f*_*mot*_ by CaD (Fig. S1).

### Flow-through chambers

The design and operation of our flow-through chambers were described before (25). Briefly, flow-through chambers (volume ∼10µL) were constructed by securing a transparent laser-cut PETG plastic coverslip to a glass coverslip coated with nitrocellulose (1.5% in amyl-acetate) using two pieces of double-sided tape. The plastic coverslip contained a through-hole under which a polycarbonate, hydrophilic, microporous membrane was placed for injection and diffusion of MLCP for myosin dephosphorylation during the assay, without creating bulk flow.

### Laser trap assay

A three-bead laser-trap assay was performed as shown in Figure 1A. Pedestals to elevate the myosin molecules were prepared by spraying 4.5μm polystyrene microspheres (171355; Polybead; Polysciences) prior to nitrocellulose coating of the glass coverslips.

Neutravidin coated trapping microspheres were prepared by incubating 3 µm amino-functionalized microspheres (171455; Polybead; Polysciences) in 8% (v/v) glutaraldehyde in 20mM PBS overnight at room temperature, pH 8. The activated microspheres were then spun down at 6000g for 6 min at 4°C and incubated with neutravidin (0.5mg/mL) and TRITC-BSA (1.25mg/mL) in 20mM PBS for 4.5h at room temperature, pH 8. The reaction was halted by washing and incubating the microspheres in ethanolamine (0.5M) for 40 min at room temperature, pH 8. The remaining non-conjugated surface of the microspheres was blocked with BSA (10mg/mL) for 30 min at 4°C and washed 10 additional times in 20mM PBS. The functionalized microspheres were stored on ice in the dark in 20mM PBS with BSA (10mg/mL).

### Laser trap assay measurements and data acquisition

Functional phosphorylated myosin (65-70ng/µL) in myosin buffer was perfused in the flow-through chamber followed by perfusion of BSA (0.5 mg/mL in myosin buffer), unlabeled filamentous actin (200 μg/mL in actin buffer) and MgATP (1 mM in actin buffer). Unbound actin filaments were flushed from the chamber with actin buffer. CaD (250nM in actin buffer) was then perfused, and the excess was washed with actin buffer. Finally, laser trap assay buffer containing the 3µm neutravidin-coated microspheres, biotinylated TRITC-labelled actin filaments, and an oxygen scavenger system (0.25 mg/ml glucose oxidase, 0.045 mg/ml catalase, and 5.75 mg/ml glucose) was added. CaD was not added to control experiments.

The flow-through chamber was then placed on the stage of an inverted microscope (Eclipse Ti; Nikon Instruments) equipped with a high-numerical-aperture oil immersion objective (CFI Plan Fluor DLL 100X, 1.3 NA; Nikon Instruments). An objective heater (Bioptechs) and a heated microscope slide holder (Chamlide TC; Quorum Technologies) were used to maintain a constant temperature of 31°C throughout the experiments. Prior to the experiment, a droplet of injection buffer was placed on top of the microporous membrane, and the ends of the flow-through chamber were sealed with vacuum grease to prevent bulk flow during MLCP injection (Fig.1A).

One neutravidin-coated microsphere was captured and held in each of the two time-shared optical traps (MMI CellManipulator Plus; Quorum Technologies) and micromanipulated to bind a biotinylated TRITC-labelled actin filament and form a bead-filament-bead dumbbell. This assembly was moved above a pedestal. Once myosin started pulling on the actin, MLCP (5μL, 3.0 μM in injection buffer) was added through the injection droplet (Fig.1A), as previously described (25).

The microspheres and actin filaments were visualized by fluorescence imaging using an excitation light source (X-Cite 120Q; Excelitas Technologies) and an electron-multiplying charge-coupled device camera (KP-E500; 720 × 480 resolution; 30 frames/s; 8-bit grayscale; Hitachi Kokusai Electric). The experiment was also recorded in bright-field using a second camera (MMI Cell Camera; 1,392 × 1,040 resolution; 30 frames/s; 24-bit RGB; Quorum Technologies), to track injection time, as seen as changes in the bright-field signal.

### Laser trap assay data and statistical analysis

The force (*F*) exerted by the myosin molecules on the actin-microsphere system was calculated using the following equation:

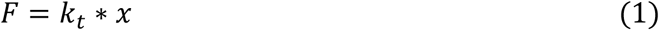

where *k*_*t*_ is the trap stiffness and *x* is the displacement of the microsphere from the center of the trap. *x* was calculated from fluorescence imaging videos using a custom Python script with the Trackpy package (38). Force events were considered if *F* exceeded 2 standard deviations above the baseline force distribution for at least 15 frames.

*T*_*inj*_ was defined as the time between the onset of the force event and the moment of MLCP injection. The frame corresponding to MLCP injection in each video was identified by monitoring changes in light intensity in the brightfield recordings using a custom Python script. Brightfield and fluorescence videos were temporally aligned by maximizing the cross-correlation of the bead-displacement traces. *T*_*Hold*_ was defined as the period between *T*_*inj*_ and the return to baseline for at least 1 sec (Fig.3A). Events where force returned to baseline due to filament breakage were not considered.

The force rise and force decay phases of force events were analyzed with an exhaustive bootstrapping approach. Specifically, we generated all unique combinations with replacement of the collected force events (126 total combinations from 5 original traces per condition) to generate distributions for force-rise and force-decay parameters. The distributions of *T*_*Hold*_ were estimated by computing the mean *T*_*Hold*_ across all force events in a given group.

For force-rise parameters, the combined traces were ensemble averaged while aligned at the start of the force event. This average trace was then fitted with the following sigmoidal function:

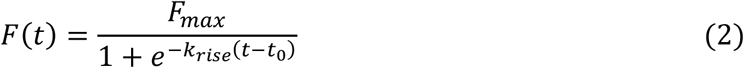

Where *t* is time in s, *F*_*max*_ is the sigmoidal upper asymptote in pN, *k*_*rise*_ is the rate of force increase in pN/s, and *t*_*O*_ is the sigmoidal midpoint in s. The fitting window extended from 10 s prior to the start of the events to the mean time of MLCP injection, calculated across all events.

For force decay parameters, the combined force events were ensemble averaged while aligned at the time of injection (t = 0). This average was then normalized to the mean force value within a 10 s window centered at 10 s after injection (t = 5 – 15 s). The data were then fitted to the following shifted-exponential decay function:

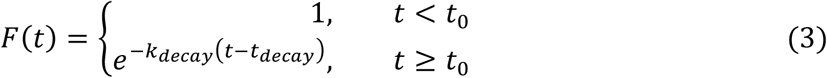

where *t* is time in s, *k*_*decay*_ is the decay rate in s^-1^, and *t*_*decay*_ is the onset of decay in s. The fitting window extended from the time of injection to 5 seconds after the mean *T*_*hol*_ calculated across all events. Fits with R^2^ values below 0.8 were excluded. Parameter distributions were compared by computing the Cliff’s *δ* effect size between conditions (39).

### IVMA

Functional phosphorylated myosin (100μg/mL) was perfused in the flow-through chamber (without pedestals), followed by perfusion of BSA (0.5 mg/mL in myosin buffer), unlabeled filamentous actin (200 μg/mL in actin buffer) and MgATP (1 mM in actin buffer). Unbound actin filaments were removed from the chamber by flushing with actin buffer. CaD (250nM in actin buffer) was then added, and the excess was washed with actin buffer. Finally, biotinylated TRITC-labelled actin (5 μg/mL) and motility assay buffer were introduced. No CaD was added for control experiments. MLCP injections were performed as described for the laser trap assay, 30s after the start of the video recording. *v*_*avg*_ and *f*_*mot*_ were measured using the motility analysis software developed by Ijpma et al. (40).

### IVMA data and statistical analysis

To assess the effects of CaD on motility decay after MLCP injection, *v*_*avg*_ and *f*_*mot*_ traces were aligned at the time of MLCP injection (t = 0). Each trace was individually normalized to a 0 - 1 scale, using the pre-injection mean as the maximum and the mean value of the last 20 s of the recording as the minimum. The resulting normalized traces were then analyzed by bootstrap-resampling (2,000 iterations) the original dataset (n = 9 per condition), ensemble averaging each sample and fitting the average trace with the sigmoidal function below:

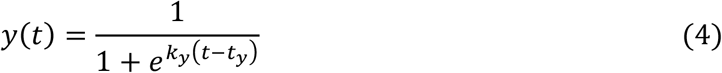

where *t* is time in s, *t*_*y*_ is the sigmoidal midpoint in s, and *k*_*y*_ is the sigmoidal rate constant in s^-1^. Fits with R^2^ values below 0.8 were excluded. Parameter distributions were compared by computing the Cliff’s *δ* effect size between conditions (39).

To determine *v*_*res*_ and *f*_*res*_ distributions, the same bootstrap-resampling procedure was applied to the non-normalized traces. After generating the ensemble averages, each trace was normalized to its pre-injection baseline, and the residual values were calculated as the mean signal during the last 20 s of the recording.

### Mixture Motility Assays

Mixture in-vitro motility assays were performed with varying ratios (0, 25, 50, 75, and 100%) of phosphorylated to unphosphorylated myosin. *v*_*avg*_ and *f*_*mot*_ data were normalized to their values measured at 100% phosphorylation. To determine the dependence of *v*_*avg*_ and *f*_*mot*_ on phosphorylation levels, the data were fitted with a linear model and a rectangular hyperbola model. The Akaike Information Criterion (41) was used to determine which model best described the data (see Supplementary Material).

The linear model was:

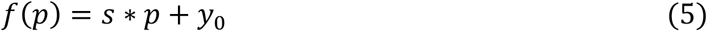

where *p* is the phosphorylation percentage, *s* is the slope of the line, and *y*_*O*_ is the Y-axis intercept.

The rectangular hyperbola model was:

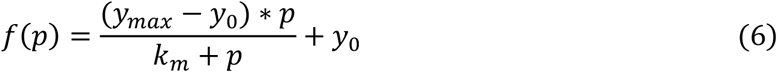

where *y*_*max*_ is the upper asymptote, *y*_*O*_ is the y-axis intercept, and *k*_*m*_ is the phosphorylation level at which *v*_*avg*_ or *f*_*mot*_ reach 50% of their maximum.

## ACKNOWLEDGEMENTS

We thank Michael Alveis (Marvid Poultry, Montreal, Quebec, Canada) for chicken gizzard donations.

## Supplementary Materials

### Force normalization by myosin molecules in proximity to the filament

To compare our observed reduction in *F*_*max*_ in the presence of CaD with the unbinding force per myosin reported by Roman et al. 2014, we normalized our median *F*_*max*_ measurements by the estimated number of myosin heads within reach of the filament (1). Following Roman et al., we assumed that myosin molecules interact with actin within a 26-nm wide band along the filament. By modelling the actin filament as a straight-line tangent to the 4.5-*μm* pedestal bead, we calculated the interacting length of the filament to be 480nm; this corresponds to the segments falling within a 13-nm reach (half of the total 26-nm band) from the pedestal surface. The surface density of myosin was estimated at ∼2000 heads/*μm*^*2*^ based on previous measurements by Warshaw and colleagues (2). The total number of proximal myosin molecules was then calculated by multiplying this density by the interaction area (26 nm x 480 nm). Dividing the measured *F*_*max*_ values by this count yielded a normalized median force of 1.02 pN/myosin in the absence of CaD. Notably, this value aligns closely with the estimated ∼1pN force derived from single-molecule experiments (3). In the presence of CaD, the normalized *F*_*max*_ was reduced to 0.54 pN/myosin.

### *T*_*Inj*_ Calculations

To ensure that the differences in *T*_*Hold*_ with and without CaD (main text Fig. 2B) were not artifacts induced by different times of injection, we calculated the time between the start of the force event and the moment of MLCP injection (*T*_*Inj*_) for each force trace collected. The *T*_*Inj*_ distribution was estimated with an exact bootstrap analysis by generating all unique combinations with replacement of the force events and calculating the mean *T*_*Inj*_ across all force events within groups. As shown in Figure S1, the *T*_*Inj*_ distributions with and without CaD had a considerable amount of overlap (Cliff’s *δ* = 0.20) with comparable median values and confidence intervals (control: 7.93 s [5.43 – 11.77] vs CaD: 7.80s [5.20 – 12.58]). Together these confirm that *T*_*Inj*_ was consistent between conditions and could not have influenced the observed *T*_*Hold*_ values.

**Figure S1:**
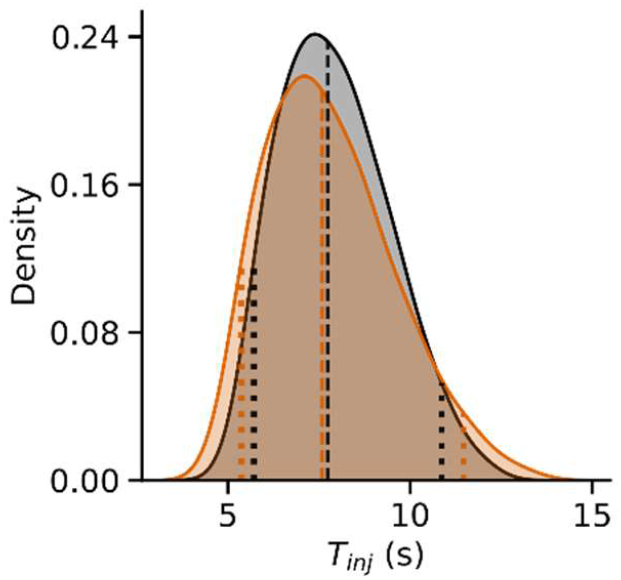
*T*_*Inj*_ distributions. Data in the presence of 250nM CaD are shown in orange; data in its absence is shown in grey. Vertical dashed lines indicate the median. Vertical dotted lines indicate the 95% CI.

### Effects Of Actin Biotinylation On Caldesmon Modulation of the IVMA

To verify whether biotinylation of actin monomers impacts the ability of CaD to regulate actomyosin interactions, we measured *v*_*avg*_ and *f*_*mot*_ in the presence and absence of 250nM CaD with biotinylated or unbiotinylated actin filaments. The data were analyzed using a two-way ANOVA followed by the Tukey test for multiple comparisons (Table S1). As shown in Figure S2A, the addition of CaD caused a comparable decrease in *v*_*avg*_ irrespective of actin biotinylation status. Similarly, CaD had no significant effect on *f*_*mot*_ for either biotinylated or unbiotinylated actin (Fig. S2B). Taken together, these results indicate that actin biotinylation does not interfere with the regulatory effect of CaD on myosin-actin mechanics.

**Table S1:** IVMA measurements with biotinylated and unbiotinylated actin.

| Actin Type | Condition | $v_{avg}$ ($\mu\text{m/s}$) | $f_{mot}$ |
| --- | --- | --- | --- |
| 0% Biotinylation | Control | $0.65 \pm 0.02$ | $0.88 \pm 0.03$ |
| | 250nM CaD | $0.50 \pm 0.03$ | $0.84 \pm 0.06$ |
| 25% Biotinylation | Control | $0.63 \pm 0.02$ | $0.86 \pm 0.04$ |
| | 250nM CaD | $0.47 \pm 0.04$ | $0.80 \pm 0.08$ |

Data are presented as mean ± SEM.

**Figure S2:**
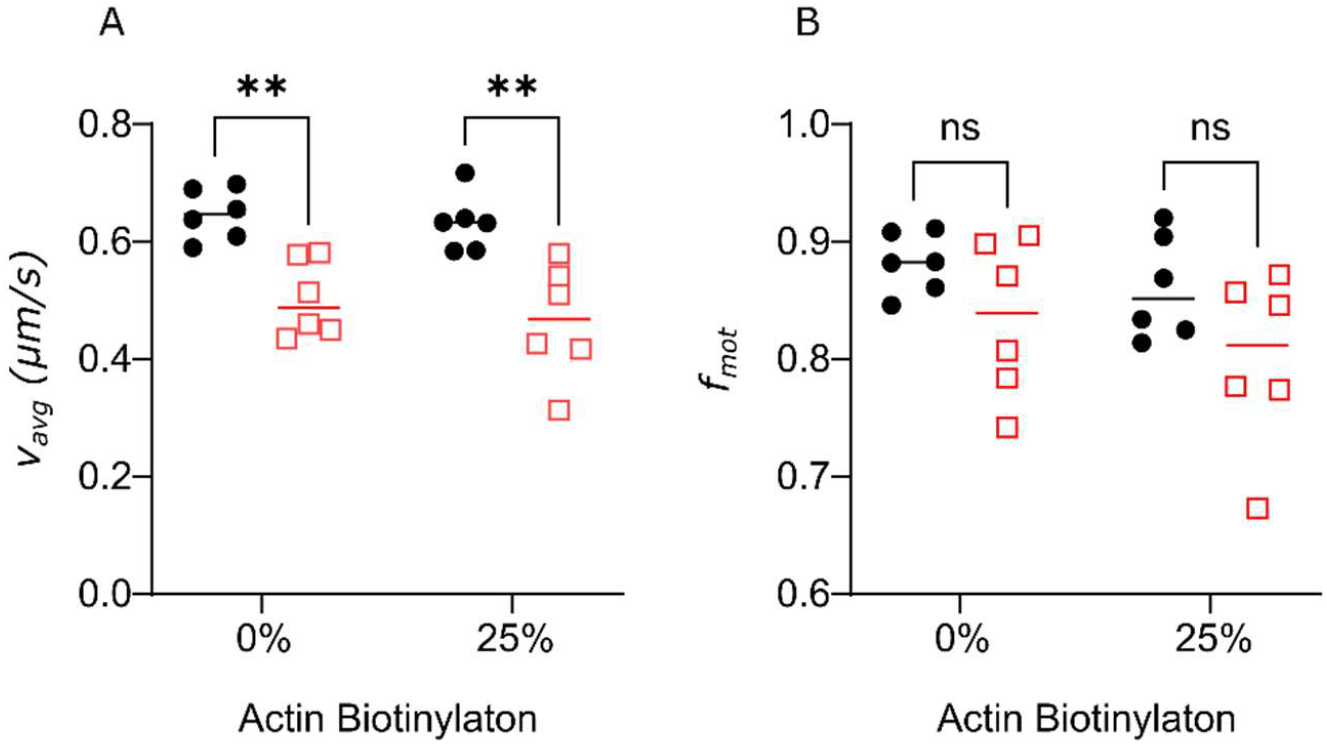
Biotinylation of actin filaments does not alter the effects of CaD on filament motility. (A) *v*_*avg*_ and (B) *f*_*mot*_ measured in the IVMA using filaments with 0% or 25% biotinylation, in the absence (black circles) or presence (red squares) of 250nM CaD. Vertical lines show the average of the individual datapoints. Groups were compared with Tukey’s HSD after an ANOVA test. **; p < 0.01

### Determination Of Model Used to Describe Mixture Motility Data

To determine whether to fit the dependence of *v*_*avg*_ and *f*_*mot*_ on phosphorylation levels with a linear or a rectangular hyperbola model, we compared both models within all datasets (*v*_*avg*_ and *f*_*mot*_ with and without CaD) using the Akaike Information Criterion (AIC)(4). All *f*_*mot*_ data and the control *v*_*avg*_ data were best described by the rectangular hyperbola. For the *v*_*avg*_ response in the presence of CaD, the data appeared quasi-linear with no observable plateau within the phosphorylation range. While the rectangular hyperbola could technically converge on this dataset, it yielded physically unrealistic parameters (projected *k*_*m*_ >> 100%). Table S2 shows the parameters of each model fit along with the *Δ*AIC comparison between models for each dataset.

**Table S2:** Comparison of mathematical models describing the phosphorylation dependence of motility parameters.

| Metric | Condition | Model | $y_{max} / s$ | $y_0$ | $k_m$ | $R^2$ | $\Delta\text{AIC}$ |
| --- | --- | --- | --- | --- | --- | --- | --- |
| $v_{avg} \mu\text{m/s}$ | Control | Rect. Hyperbola | $0.8 \pm 0.1$ | $0.20 \pm 0.04$ | $37 \pm 17$ | 0.83 | 13.86 |
| | | Linear | $4.2 \pm 0.6 \times 10^{-3}$ | $0.29 \pm 0.04$ | --- | 0.69 | |
| | 250nM CaD | Rect. Hyperbola | $2.0 \pm 1.4$ | $0.10 \pm 0.02$ | $312 \pm 17$ | 0.91 | 0.25 |
| | | Linear | $4.6 \pm 0.3 \times 10^{-3}$ | $0.12 \pm 0.02$ | --- | 0.90 | |
| $f_{mot}$ | Control | Rect. Hyperbola | $1.00 \pm 0.05$ | $0.52 \pm 0.03$ | $12 \pm 6$ | 0.85 | 25.09 |
| | | Linear | $3.7 \pm 0.6 \times 10^{-3}$ | $0.65 \pm 0.04$ | --- | 0.58 | |
| | 250nM CaD | Rect. Hyperbola | $1.00 \pm 0.07$ | $0.23 \pm 0.04$ | $18 \pm 6$ | 0.92 | 17.03 |
| | | Linear | $6.9 \pm 0.6 \times 10^{-3}$ | $0.35 \pm 0.04$ | --- | 0.83 | |

Mixture motility data for *v*_*avg*_ and *f*_*mot*_ with and without 250nM CaD were fitted to two candidate models: a rectangular hyperbola and a linear model. Values are reported as best fit ± standard error. The column *y*_*max*_*/s* reports the projected asymptotic maximum (*y*_*max*_) for the hyperbolic fit or the slope (*s*) for the linear fit. *y*_*O*_ represents the y-intercept (baseline activity at 0% phosphorylation), and *k*_*m*_ denotes the phosphorylation level at half-maximal response derived form the hyperbolic fit. *ΔAIC* indicates the magnitude of the difference in AIC between the two models (*ΔAIC = /AIC*_*linear*_ − *AIC*_*hyperbola*_*/*). A *ΔAIC < 2*, as observed for the *v*_*avg*_ in the presence of 250nM CaD, indicates that the rectangular hyperbola offers no statistical advantage over the simpler linear model. Parameters are presented as best-fit values ± SE.

### Analytical estimation of dephosphorylation kinetics

#### Mathematical Framework

The rate of myosin dephosphorylation 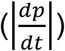 was expressed as the product of the observable *v*_*avg*_ decay rate 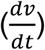 during myosin dephosphorylation and the inverse sensitivity of *v*_*avg*_ to phosphorylation levels 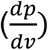:

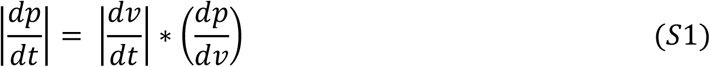

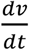 was obtained by differentiating a sigmoidal function fitted to the non-normalized *v*_*avg*_ data during MLCP-induced myosin dephosphorylation presented in the main text (Fig. 3A). Note that, unlike the analysis in the main text (which used normalized data to compare relative rates), here the estimation used non-normalized traces. Hence, the *v*_*avg*_ data was fitted with the function below:

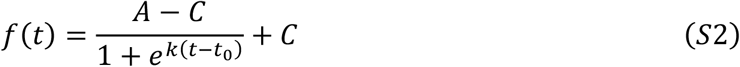

where A and C are the upper and lower sigmoidal plateaus respectively, *k* is the sigmoidal rate, and *t*_*O*_ is the sigmoidal midpoint.

Differentiating Eq. S2 to obtain 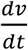 yields the function below:

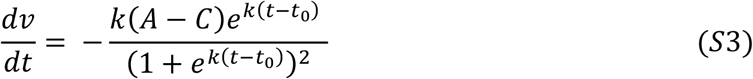

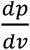 was calculated as the reciprocal of the derivative of the function fitted to the *v*_*avg*_ vs myosin phosphorylation data presented in the main text (Eqs. 4 & 5 and Fig. 4A). For conditions containing 250nM CaD, which were fitted with a linear model, 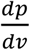 was defined as:

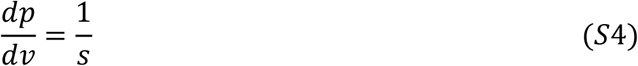

For control condition, which were fitted with a rectangular hyperbola model, 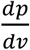 was defined as:

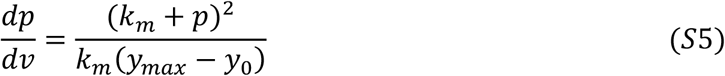

Because the sensitivity in Eq. S5 depends on the instantaneous phosphorylation *p*, we solved for *p*(*t*) at each time point by inverting the rectangular hyperbola (Main Text Eq. 5) and inputting the instantaneous *v*_*avg*_ during MLCP injection (Eq. S2):

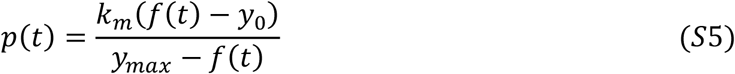

where *f(t)* is the instantaneous *v*_*avg*_ during MLCP injection described by the sigmoidal function in Eq. S2. This derived *p*(*t*) was then substituted into Eq. S4 to determine 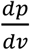.

#### Error Propagation

The variation in *v*_*avg*_ decay during MLCP injection was captured by bootstrap resampling the *v*_*avg*_ traces 2000 times. Each resampled average trace was fitted with Eq. S2, yielding a distribution of 2000 sets of time-course parameters (*A, C, k, t*_*O*_) that reflect the experimental variability.

To account for the statistical error in *v*_*avg*_ vs myosin phosphorylation models (Main text Eq. 3 or Eq.4), we generated 2000 synthetic calibration curves by sampling the calibration parameters ({*y*_*max*_, *y*_*O*_, *k*_*m*_*} or {s, y*_*O*_}) from a multivariate normal distribution defined by the optimal best-fit parameters and their covariance matrix obtained from the non-linear least squares optimization.

We calculated the dephosphorylation rate trajectory by pairing the *i*-th bootstrapped time course parameters with the *i*-th sampled calibration parameters. The reported rates represent the mean of these simulations, with 95% confidence intervals defined by the 2.5^th^ and the 97.5^th^ percentiles.

#### Results

To verify whether the presence of CaD significantly altered the rate of myosin dephosphorylation, we compared the distribution of the estimated dephosphorylation rates with and without CaD. We calculated the time-dependent difference between the conditions for each simulation iteration as:

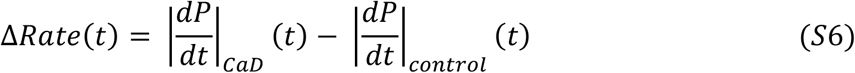

From these differential traces, we constructed a 95% confidence interval for the difference *ΔRate(t)*. We defined a statistically significant effect as any time interval where this 95% confidence interval did not include zero.

The results of this analytical estimation for the *v*_*avg*_ data are shown in Fig. S3. The derived dephosphorylation rates (Fig. S3B) revealed a substantial overlap in dephosphorylation rates between both conditions (control and CaD). In both the control and CaD groups, the dephosphorylation rate followed a bell-shaped curve: increasing gradually after MLCP injection, peaking mid-way through dephosphorylation, and declining thereafter.

Crucially, the 95% confidence interval for *ΔRate* included zero across the entire time-course (Fig. S3D). This indicates that the presence of CaD did not significantly alter the enzymatic rate of myosin dephosphorylation. Notably, despite observing no statistical difference in the instantaneous rates, the absolute levels of myosin phosphorylation were significantly higher in the presence of CaD for a brief interval between 16.7 s and 37.3 s post-injection (Fig. S3C). However, the lower bound of the confidence interval remained close to zero during this period.

**Figure S3:**
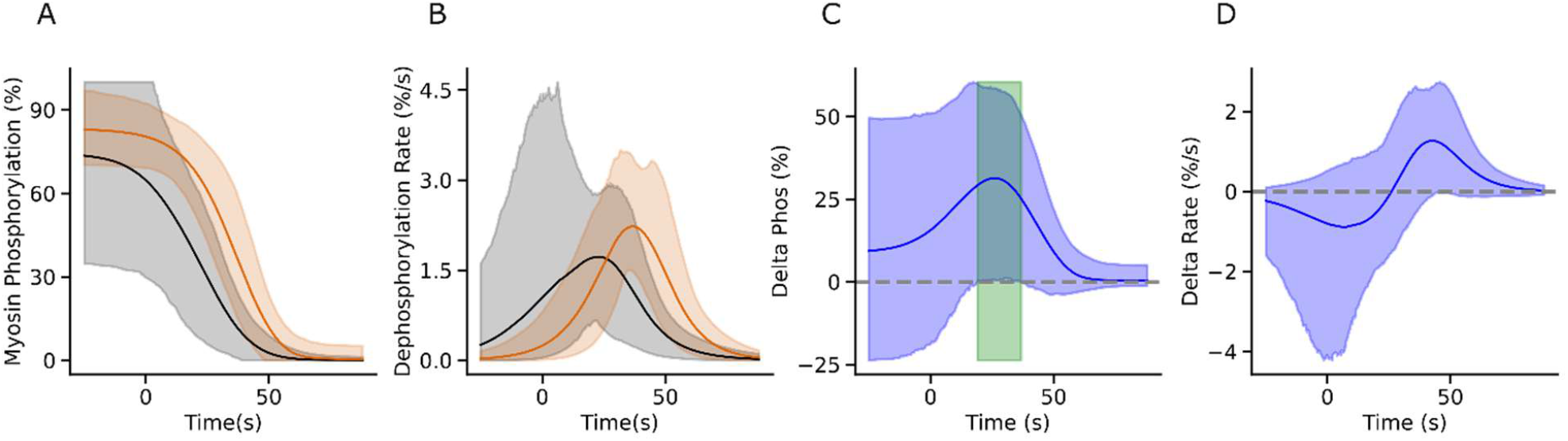
Analytical Estimation of LC_20_ dephosphorylation kinetics from *v*_*avg*_ decay data. (A) Estimated time-course of myosin phosphorylation levels (*p*(*t*)). (B) Estimated dephosphorylation rates 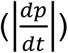. (C) The difference in estimated phosphorylation levels (*ΔP)*. Green shading highlights the interval (16.7–37.3 s) where phosphorylation levels were significantly higher in the presence of CaD (95% CI > 0). (D) The time-dependent difference in dephosphorylation rates (*ΔRate)*. Solid lines represent the average; shaded regions indicate 95% confidence intervals.

## REFERENCES

1. P. F. Dillon, M. O. Aksoy, S. P. Driska, R. A. Murphy, Myosin Phosphorylation and the Cross-Bridge Cycle in Arterial Smooth Muscle. Science 211, 495–497 (1981).

2. P. F. Dillon, R. A. Murphy, High force development and crossbridge attachment in smooth muscle from swine carotid arteries. Circulation Research 50, 799–804 (1982).

3. M. g. Tansey, M. Hori, H. Karaki, K. e. Kamm, J. t. Stull, Okadaic acid uncouples myosin light chain phosphorylation and tension in smooth muscle. FEBS Letters 270, 219–221 (1990).

4. M. O. Aksoy, S. Mras, K. E. Kamm, R. A. Murphy, Ca2+, cAMP, and changes in myosin phosphorylation during contraction of smooth muscle. American Journal of Physiology-Cell Physiology 245, C255–C270 (1983).

5. W. Fischer, G. Pfitzer, Rapid myosin phosphorylation transients in phasic contractions in chicken gizzard smooth muscle. FEBS Letters 258, 59–62 (1989).

6. K. Y. Horiuchi, S. Chacko, Caldesmon inhibits the cooperative turning-on of the smooth muscle heavy meromyosin by tropomyosin-actin. Biochemistry 28, 9111–9116 (1989).

7. M. E. Hemric, J. M. Chalovich, Effect of caldesmon on the ATPase activity and the binding of smooth and skeletal myosin subfragments to actin. Journal of Biological Chemistry 263, 1878–1885 (1988).

8. C.-L. Albert Wang, Caldesmon and smooth-muscle regulation. Cell Biochem Biophys 35, 275–288 (2001).

9. J. A. Lash, J. R. Sellers, D. R. Hathaway, The Effects of caldesmon on smooth muscle heavy actomeromyosin ATPase activity and binding of heavy meromyosin to actin. Journal of Biological Chemistry 261, 16155–16160 (1986).

10. J. R. Haeberle, K. M. Trybus, M. E. Hemric, D. M. Warshaw, The Effects of smooth muscle caldesmon on actin filament motility. Journal of Biological Chemistry 267, 23001–23006 (1992).

11. H. N. Roman, et al., The Role of Caldesmon and its Phosphorylation by ERK on the Binding Force of Unphosphorylated Myosin to Actin. Biochim Biophys Acta 1840, 3218–3225 (2014).

12. K. Mabuchi, Y. Li, A. Carlos, C.-L. A. Wang, P. Gracefa, Caldesmon exhibits a clustered distribution along individual chicken gizzard native thin filaments. J Muscle Res Cell Motil 22, 77–90 (2001).

13. S. B. Kokate, et al., Caldesmon controls stress fiber force-balance through dynamic cross-linking of myosin II and actin-tropomyosin filaments. Nat Commun 13, 6032 (2022).

14. K. Albrecht, A. Schneider, C. Liebetrau, J. C. Rüegg, G. Pfitzer, Exogenous caldesmon promotes relaxation of guinea-pig skinned taenia coli smooth muscles: inhibition of cooperative reattachment of latch bridges? Pflügers Arch 434, 534–542 (1997).

15. J. J. Earley, X. Su, R. S. Moreland, Caldesmon Inhibits Active Crossbridges in Unstimulated Vascular Smooth Muscle. Circulation Research 83, 661–667 (1998).

16. S. Pütz, et al., Caldesmon ablation in mice causes umbilical herniation and alters contractility of fetal urinary bladder smooth muscle. Journal of General Physiology 153, e202012776 (2021).

17. H. Guo, et al., Ablation of Smooth Muscle Caldesmon Afects the Relaxation Kinetics of Arterial Muscle. Pflugers Arch 465, 283–294 (2013).

18. G. Pfitzer, C. Zeugner, M. Troschka, J. M. Chalovich, Caldesmon and a 20-kDa actin-binding fragment of caldesmon inhibit tension development in skinned gizzard muscle fiber bundles. Proceedings of the National Academy of Sciences 90, 5904–5908 (1993).

19. U. Malmqvist, A. Arner, R. Makuch, R. Dabrowska, The Effects of caldesmon extraction on mechanical properties of skinned smooth muscle fibre preparations. Pflugers Arch. 432, 241–247 (1996).

20. E. M. Smolock, et al., siRNA-mediated knockdown of h-caldesmon in vascular smooth muscle. American Journal of Physiology-Heart and Circulatory Physiology 297, H1930– H1939 (2009).

21. T. Okagaki, S. Higashi-Fujime, R. Ishikawa, H. Takano-Ohmuro, K. Kohama, In Vitro Movement of Actin Filaments on Gizzard Smooth Muscle Myosin: Requirement of Phosphorylation of Myosin Light Chain and Effects of Tropomyosin and Caldesmon1. The Journal of Biochemistry 109, 858–866 (1991).

22. V. P. Shirinsky, K. G. Biryukov, J. M. Hettasch, J. R. Sellers, Inhibition of the relative movement of actin and myosin by caldesmon and calponin. Journal of Biological Chemistry 267, 15886–15892 (1992).

23. K. Y. Horiuchi, S. Chacko, Effect of unphosphorylated smooth muscle myosin on caldesmon-mediated regulation of actin filament velocity. J Muscle Res Cell Motil 16, 11–19 (1995).

24. I. D. C. Fraser, S. B. Marston, In Vitro Motility Analysis of Smooth Muscle Caldesmon Control of Actin-Tropomyosin Filament Movement (*). Journal of Biological Chemistry 270, 19688–19693 (1995).

25. M. J. Hammell, L. Kachmar, Z. Balassy, G. IJpma, A.-M. Lauzon, Molecular-level evidence of force maintenance by smooth muscle myosin during LC20 dephosphorylation. Journal of General Physiology 154, e202213117 (2022).

26. A. Sen, Y.-D. Chen, B. Yan, J. M. Chalovich, Caldesmon Reduces the Apparent Rate of Binding of Myosin S1 to Actin—Tropomyosin. Biochemistry 40, 5757–5764 (2001).

27. L. Velaz, Y. D. Chen, J. M. Chalovich, Characterization of a caldesmon fragment that competes with myosin-ATP binding to actin. Biophysical Journal 65, 892–898 (1993).

28. B. Yan, A. Sen, J. M. Chalovich, Y. Chen, Theoretical Studies on Competitive Binding of Caldesmon and Myosin S1 to Actin: Prediction of Apparent Cooperativity in Equilibrium and Slow-Down in Kinetics of S1 Binding by Caldesmon. Biochemistry 42, 4208–4216 (2003).

29. S. Marston, K. Pinter, P. Bennett, Caldesmon binds to smooth muscle myosin and myosin rod and crosslinks thick filaments to actin filaments. J Muscle Res Cell Motil 13, 206–218 (1992).

30. S. Chacko, E. Eisenberg, Cooperativity of actin-activated ATPase of gizzard heavy meromyosin in the presence of gizzard tropomyosin. Journal of Biological Chemistry 265, 2105–2110 (1990).

31. G. Pfitzer, et al., Is Myosin Phosphorylation Suficient to Regulate Smooth Muscle Contraction? in Sliding Filament Mechanism in Muscle Contraction, H. Sugi, Ed. (Springer US, 2005), pp. 319–328.

32. Y. Li, et al., The Major Myosin-binding Site of Caldesmon Resides Near Its N-terminal Extreme*. Journal of Biological Chemistry 275, 10989–10994 (2000).

33. M. E. Hemric, J. M. Chalovich, Characterization of Caldesmon Binding to Myosin. J Biol Chem 265, 19672–19678 (1990).

34. K. M. Trybus, Biochemical Studies of Myosin. Methods 22, 327–335 (2000).

35. A. Sobieszek, “Smooth Muscle Myosin: Molecule Conformation, Filament Assembly and Associated Regulatory Enzymes” in Airways Smooth Muscle: Biochemical Control of Contraction and Relaxation, D. Raeburn, M. A. Giembycz, Eds. (Birkhäuser, 1994), pp. 1–29.

36. A. Sobieszek, J. Borkowski, V. S. Babiychuk, Purification and Characterization of a Smooth Muscle Myosin Light Chain Kinase-Phosphatase Complex*. Journal of Biological Chemistry 272, 7034–7041 (1997).

37. A. Sobieszek, E. B. Babiychuk, B. Ortner, J. Borkowski, Purification and Characterization of a Kinase-associated, Myofibrillar Smooth Muscle Myosin Light Chain Phosphatase Possessing a Calmodulin-targeting Subunit*. Journal of Biological Chemistry 272, 7027–7033 (1997).

38. D. B. Allan, T. Caswell, N. C. Keim, C. M. van der Wel, trackpy: Trackpy v0.4.1. (2018). 10.5281/zenodo.1226458. Deposited 21 April 2018.

39. N. Clif, Dominance statistics: Ordinal analyses to answer ordinal questions. Psychological Bulletin 114, 494–509 (1993).

40. G. Ijpma, Z. Balassy, A.-M. Lauzon, Rapid time-stamped analysis of filament motility. J Muscle Res Cell Motil 39, 153–162 (2018).

41. H. Akaike, A new look at the statistical model identification. IEEE Transactions on Automatic Control 19, 716–723 (1974).

## References to the Supplementary Materials

1. H. N. Roman, et al., The Role of Caldesmon and its Phosphorylation by ERK on the Binding Force of Unphosphorylated Myosin to Actin. Biochim Biophys Acta 1840, 3218–3225 (2014).

2. D. E. Harris, D. M. Warshaw, Smooth and skeletal muscle myosin both exhibit low duty cycles at zero load in vitro. Journal of Biological Chemistry 268, 14764–14768 (1993).

3. A. M. Lauzon, et al., A 7-amino-acid insert in the heavy chain nucleotide binding loop alters the kinetics of smooth muscle myosin in the laser trap. J Muscle Res Cell Motil 19, 825–837 (1998).

4. H. Akaike, A new look at the statistical model identification. IEEE Transactions on Automatic Control 19, 716–723 (1974).

